# Exposure to the antimicrobial peptides LL-37 and ATRA-1 induces a lipidome response in *Staphylococcus aureus* that alters membrane biophysical properties

**DOI:** 10.64898/2026.04.16.718754

**Authors:** Christian Fuertes-Chaves, Juan E. Gonzalez, Elizabeth Suesca, Paula Guzmán-Sastoque, Carolina Muñoz-Camargo, Marcela Manrique-Moreno, Chiara Carazzone, Chad Leidy

## Abstract

*Staphylococcus aureus* (*S. aureus*) is an opportunistic pathogen that is a global health concern for its ability to cause a wide spectrum of clinical infections. Due to the emergence of resistance to commonly used antibiotics, there has been interest in exploring the use of antimicrobial peptides to treat *S. aureus* infections. However, changes in the lipid composition of the lipid bilayer membrane can alter the activity of peptides, and *S. aureus* is able to induce variations in lipid composition in response to environmental stress. Here, we explore how the main lipid components in *S. aureus* are altered when exposed to LL-37, a human cathelicidin involved in primary immune response, and ATRA-1, a short antimicrobial peptide derived from the snake *Naja atra* venom. A lipidomic study is conducted through HPLC-MS-MS (LC-ESI-MS/MS) to quantify phosphatidylglycerol, cardiolipin, lysyl-phosphatidylglycerol, monogalacto- and digalacto-diacylglycerol, and carotenoids. In addition, menaquinones, responsible for electron transport during oxidative phosphorylation, were also quantified. Biophysical properties such as membrane electric surface potential and lipid packing were assessed. We find that lipid adaptation is specific to the type of antimicrobial peptide, where ATRA-1 mainly induces changes in the electric surface potential through variations in Lysyl-PG, while exposure to LL-37 changes carotenoid levels, inducing an increase in membrane rigidity as measured by FTIR. In addition, both peptides induce a reduction in menaquinone and DGDG levels. These findings highlight the role of membrane lipid remodeling as a peptide-specific response mechanism in *S. aureus*, with implications for the development of AMP-based therapies.

**Highlights:**

- *Staphylococcus aureus* responds through shifts in lipid composition and membrane biophysical properties to exposure to the antimicrobial peptides LL-37 and ATRA-1.
- Both LL-37 and ATRA-1 lead to shifts in the glycolipids MGDG and DGDG; two lipids involved in regulating negative membrane curvature stress and responsible for shifting resistance to antimicrobial peptide activity in *Staphylococcus aureus*.
- LL-37 treatment leads to an overall reduction in carotenoid content in *Staphylococcus aureus,* including the carotenoid end-product staphyloxanthin and the precursor 4,4’-diaponeurosporenoic acid. Both lipids regulate membrane biophysical properties and protect *Staphylococcus aureus* from oxidative stress.
- Both LL-37 and ATRA-1 lead to a reduction in menaquinone levels, which are involved in the electron transport chain during oxidative phosphorylation. Reduction in these menaquinones have been associated to the formation of small colony variants that are often observed in chronic *Staphylococcus aureus* infections.

## 1. Introduction

Among the various antibiotic-resistant pathogens that pose a global public health concern, *Staphylococcus aureus* (*S. aureus*) is an opportunistic pathogen associated with significant morbidity and mortality worldwide, causing 20-30% of bloodstream and surgical site infections [1]. *S. aureus* bacteremia incidence ranges from 4.3-38.2 per 100,000 person-years with mortality levels ranging from 10-40% [2–4]. *S. aureus* is also an opportunistic commensal microorganism, with approximately 20-30% of adults colonized by this microorganism in their epithelial or mucosal tissues, where 1 to 3% of adults are colonized by methicillin-resistant *Staphylococcus aureus* (MRSA) [5, 6]. *S. aureus* can infect a variety of tissues, leading to conditions such as infectious endocarditis, osteoarticular infections, skin and soft tissue infections, pleuropulmonary infections, and widespread bacteremia [7].

*S. aureus* is an exceptionally adaptive species, capable of surviving on abiotic surfaces such as plastic, fabric, and metal for up to 90 days [8]. Consequently, it has become one of the most common types of nosocomial (hospital-acquired) infections [9]. Antibiotic resistance in *S. aureus* arises through chromosomal mutations, plasmid-mediated acquisition of resistance genes, biofilm formation, and the emergence of persister cells, which are genetically similar but phenotypically distinct dormant subpopulations capable of surviving antibiotic stress until favorable conditions return [10].

A widespread primary defense mechanism against bacterial infections in multicellular organisms is the biosynthesis of cationic antimicrobial peptides (AMPs). These short peptides, typically ranging from 10 to 40 amino acids in length, selectively bind to negatively charged bacterial membranes through non-specific electrostatic interactions [11]. Unlike conventional antibiotics that rely on specific protein targets or receptors, AMPs exert bactericidal activity by directly interacting with membrane phospholipids, disrupting bilayer integrity, forming pores, and inducing membrane depolarization [12, 13].

In humans, three major AMP groups contribute to innate immunity: defensins, histatins, and cathelicidins [14]. Among these, LL-37 is the only human cathelicidin-derived peptide that has demonstrated significant anti-staphylococcal activity [15, 16]. LL-37 is a 37-amino-acid peptide that adopts an α-helical structure upon interaction with anionic membranes and carries a net positive charge of +6 under physiological conditions. One proposed mode of action involves the formation of toroidal tetrameric transmembrane pores, leading to disruption of the electrochemical gradient [17–19]. LL-37 also plays a role in modulating the immune response in humans [20].

Another AMP of interest is ATRA-1, a short cationic peptide derived from the Chinese cobra (*Naja atra*) cathelicidin. Although not part of human physiology, ATRA-1 exhibits strong anti-biofilm activity and antimicrobial activity against *S. aureus*, with its full range of properties still under investigation [21, 22]. ATRA-1 has an α-helical structure and carries a net positive charge of +8 [23]. Due to its relatively short length, ATRA-1’s primary mechanism of action is carpet-like or detergent-like disruption of the membrane [21]. Importantly, the pore-forming or disruptive activity of both LL-37 and ATRA-1 is highly sensitive to membrane lipid composition [21].

The membrane of *S. aureus* is predominantly composed of the anionic phospholipid phosphatidylglycerol (PG), which can incorporate two additional chains by reacting with phosphatidic acid to form the tetra-acyl-chain phospholipid cardiolipin (CL) [24]. PG can also be enzymatically modified via esterification with L-lysine to generate lysyl-phosphatidylglycerol (Lysyl-PG), a cationic phospholipid that reduces the binding affinity of AMPs to the bacterial surface [25]. Notably, *S. aureus* exhibits particularly high levels of Lysyl-PG compared to many other bacterial species [26]. In response to environmental stressors and host defenses, including AMP exposure, *S. aureus* modulates the relative abundance of CL [27, 28] and Lysyl-PG [29, 30].

Beyond phospholipids, *S. aureus* synthesizes carotenoids [31–33], including 34 distinct variants, among which staphyloxanthin (STX) is the most prominent [33]. STX provides the bacterium with its characteristic yellow-golden pigmentation and acts as a potent antioxidant [34–37]. Additionally, STX contributes to membrane mechanical stability. The conjugated double-bond triterpenoid of STX restricts molecular conformational freedom, rendering the molecule more rigid and compact than typical phospholipids [32, 38].

*S. aureus* membranes also contain the uncharged galactolipids monogalactodiacylglycerol (MGDG) and digalactodiacylglycerol (DGDG) [39, 40]. MGDG is characterized for having a conical shape that induces negative curvature stress in the membrane and promotes the non-lamellar inverted hexagonal phase. A reduced DGDG/MGDG ratio leads to more accentuated membrane curvature stress. Recent results also suggest that, when *S. aureus* incorporates unsaturated lipids from the growth media [39], the conicity of MGDG and DGDG increases due to incorporation of oleic acid into these glycolipids. This leads to accentuated negative curvature stress in *S. aureus* membranes. This effect has been shown to reduce the activity of antimicrobial peptides [39].

Menaquinones (MKs) are also essential membrane components in *S. aureus*, functioning as electron carriers in the electron transport chain under both aerobic and anaerobic conditions and contributing to virulence, cytochrome formation, and broader metabolic regulation [41, 42]. Alterations in MK biosynthesis may reflect adaptive metabolic responses to AMP-induced stress, and modulation of MK levels has been associated with mitigation of haem-related oxidative stress [43].

In this study, we investigate how the membrane lipid composition and biophysical properties of *S. aureus* are modulated in response to exposure to sublethal concentrations of LL-37 and ATRA-1. We examine shifts in phospholipid composition, including changes in the relative abundance of PG, CL, and Lysyl-PG, MGDG, and DGDG, as well as variations in carotenoid and menaquinone content. These compositional changes are specific to peptide treatment and correlate with alterations in membrane biophysical properties, including lipid phase behavior measured by Fourier transform infrared (FTIR) spectroscopy, and membrane surface charge density assessed via zeta potential. The lipid phase behavior of *S. aureus* has been shown to modulate antimicrobial activity, in particular for the human phospholipase PLA2 type IIA [44]. Understanding how *S. aureus* dynamically remodels its membrane in response to antimicrobial peptides may facilitate the identification of lipid biosynthetic pathways that contribute to adaptive resistance and represent potential therapeutic targets.

## 2. Materials and Methods

### 2.1. Reagents

Analytical-grade methanol (MeOH) and chloroform (CHCl3) were purchased from Honeywell (Michigan, USA). 2,6-Di-tert-butyl-4-methylphenol 0.1 % (DTBH), powder components for Luria-Bertani medium (Tryptone, Yeast and NaCl) and HEPES buffer were purchased from Sigma-Aldrich (St. Louis, MO, USA). HPLC-grade water was obtained from a water purification system Heal Force Smart-Mini (Shanghai, China). Peptides LL-37 (LLGDFFRKSKEKIGKEFKRIVQRIKDFLRNLVPRTES) and ATRA-1 (KRFKKFFKKLK-NH2) were purchased from GenScript Biotech (New Jersey, USA).

### 2.2. Bacterial Cultures

The *S. aureus* strain used in this study is cataloged as SA401 and was provided as a clinical isolate by the Microbiological Research Center of the Universidad de Los Andes (CIMIC). SA401 was retrieved from stock stored at -80 °C.

Under sterile conditions, using the tip of a micropipette, a portion of the inoculum was streaked onto Petri dishes containing Luria-Bertani (LB) agar medium (10 g of tryptone, 10 g of NaCl and 5 g of yeast extract per 15 g of agar). The plate was incubated at 37 °C for at least 12 hours to obtain visible individual colonies. After this period, the plate was sealed with Parafilm and stored in a refrigerator at 4 °C, remaining suitable for colony extraction for about a month. After this time, a new plate had to be prepared.

For each sample, a single colony was taken and inoculated into 10 ml of LB medium with constant agitation at 150 rpm and 37 °C for 18 hours. Then, an aliquot of 10 μl was seeded in 150 ml of fresh LB, in 500 ml capacity Erlenmeyer flasks for optimal oxygenation conditions, at the same time the peptide was added at sub-lethal concentrations in the treated samples, resulting in concentrations of 40 µg/ml and 30 µg/ml for the LL-37 and ATRA-1 samples respectively. The new growth was incubated for 24 hours into a shaker at 150 rpm and 37 °C.

### 2.3. Zeta potential measurements

To obtain reliable measurements of the membrane surface electric potential, unilamellar liposomes with a diameter of 100 nm were formed in HEPES buffer (20mM + NaCl 180 mM) via extrusion from *S. aureus* extracted lipids. Instrument parameters were set to a viscosity of 0.8882 cP and a refractive index of 1.33. The refraction index of samples was taken as 1.45 and the absorbance as 0.001. To ensure a size distribution of vesicles centered around 100 nm, particle size was measured by Dynamic Light Scattering (DLS) and peak values of particle diameter were obtained; after verifying adequate sizes of liposomes, zeta Potential was measured 4 times per sample, each measurement consisted of 100 runs. Both measurements were made using ZETASIZER Nano Series ZSP (Malvern Instruments).

The surface charge density (𝜎) of the liposomes was calculated from the measured Zeta potential (𝜁) using the Grahame equation [45], accounting for the non-linear distribution of ions within the electrical double layer at high ionic strengths for a 1:1 electrolyte [46]:

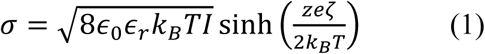

Where 𝜖_0_is the permittivity of vacuum (8.854 × 10^−12^ F/m), 𝜖_𝑟_ is the relative permittivity of the medium (∼78.5 for water), 𝑘_𝐵_ is the Boltzmann constant (1.381 × 10^−23^ J/K), T is the absolute temperature (298.15 K), and 𝐼 is the ionic strength of the bulk solution (calculated as 190 mM, accounting for the 180 mM NaCl and the ionic contribution of the 20 mM HEPES buffer at pH 7.4. For use in the Grahame Equation, this was converted to ions/m^3^ using the relation 𝐼 ⋅ 1000 ⋅ 𝑁_𝐴_ (where 𝑁_𝐴_is Avogadro’s number). **z** is the valence of the electrolyte (**z**=1 for NaCl), **e** is the elementary charge, and **ζ** is the measured zeta potential in Volts.

### 2.4. Scanning Electron Microscopy (SEM)

Cell morphologies were examined by SEM before and after exposure to the peptide at 30 kx and 200 kx magnifications with an accelerating voltage of 10 kV. Cells were harvested by centrifugation of 1 mL of overnight culture to obtain a cell pellet. All samples were fixed in 2.5% glutaraldehyde for 24 h, rinsed with HPLC-grade water to remove salts, and dehydrated through a graded ethanol series (70%, 95%, and 100% v/v, 30 min each). Samples were then sputter-coated with gold prior to imaging.

### 2.5. Separation of total lipids

Total lipids were extracted following a modified Bligh and Dyer–type protocol adapted for bacterial cells and carotenoid preservation. This extraction protocol was selected to ensure efficient recovery of both polar membrane lipids and hydrophobic isoprenoids such as carotenoids and menaquinones. Briefly, cell suspensions were centrifuged at 8500 rpm, and the resulting pellets were snap-frozen in liquid nitrogen and lyophilized for 24 h. An aliquot corresponding to 200 mg of dry cell weight was transferred to glass tubes containing 10 glass beads and mechanically disrupted. Samples were extracted with 5 mL of MeOH:CHCl3 (2:1, v/v) supplemented with 0.1% (w/v) 2,6-di-tert-butyl-4-methylphenol (DTBH) to prevent oxidation of carotenoids and unsaturated lipids. The mixtures were vortexed for 5 min. Subsequently, 1.5 mL of 1.7 M NaCl solution was added to induce phase separation, and samples were vortexed for 1 min every 15 min over a total extraction period of 2 h. An additional 5 ml of MeOH:CHCl3 (2:1, v/v) was then added, followed by vortexing for 5 min and the addition of 1.5 ml of 1.7 M NaCl. Samples were again mixed intermittently (1 min every 15 min for 2 h) and centrifuged at 5000 rpm for 15 min at 4 °C to facilitate phase separation. The extended, two-cycle extraction (total 4 h) was optimized to maximize recovery of both polar and non-polar lipid classes while minimizing oxidative degradation through the use of BHT, low temperature, and limited light exposure. The lower chloroform phase, containing the extracted lipids, was carefully collected using a separation funnel.

To maximize lipid recovery, the residual pellet was re-extracted with 10 mL of MeOH:CHCl3 (2:1, v/v), vortexed for 5 min, followed by the addition of 3 mL of 1.7 M NaCl and further mixing for 5 min. After centrifugation under the same conditions, the chloroform phase was collected and pooled with the previous extracts. A washing step was performed by adding methanol (half of the chloroform volume) and an equal volume of NaCl solution to the pooled organic phase. The mixture was gently agitated, periodically vented, and allowed to stand overnight in the dark to ensure complete phase separation and removal of non-lipid contaminants. The final chloroform phase was recovered, dried under a stream of nitrogen, and further desiccated by lyophilization for at least 6 h in pre-weighed amber vials.

### 2.6. Semi-targeted lipidomics by LC–MS/MS

Semi-targeted lipidomic analysis focused on five lipid classes relevant to *S. aureus* membrane biology: phosphatidylglycerols (PG), lysyl-phosphatidylglycerols (Lysyl-PG), cardiolipins (CL), monoglycosyldiacylglycerols (MGDG), and diglycosyldiacylglycerols (DGDG). Analyses were performed using a Dionex UltiMate 3000 UHPLC system (Thermo Scientific, San Jose,CA, USA) equipped with an online degasser, binary pump, autosampler, and thermostated column compartment, coupled to an LCQ Fleet ion trap mass spectrometer via an electrospray ionization (ESI) source operating in both positive and negative modes. Data acquisition and processing were carried out using Xcalibur 4.3 software. Dried lipid extracts were reconstituted at 1000 ppm in CHCl3:MeOH (1:2, v/v). A 3 μL injection volume was used, with samples maintained at 15 °C in the autosampler. Chromatographic separation was achieved on a Kinetex C18 column (2.1 × 50 mm, 2.6 μm; Phenomenex) protected by a C18 guard cartridge (4 × 2 mm, 3 μm), maintained at 50 °C. The mobile phases consisted of (A) acetonitrile with 10 mM ammonium formate (6:4, v/v) and (B) 2-propanol:acetonitrile (9:1, v/v). The flow rate was 300 μL min⁻¹. The gradient program was as follows: 45% B (0–2 min), linear increase to 65% B (2–20 min), increase to 100% B (20–31 min), held at 100% B (31–35 min), returned to 45% B (35–37 min), and equilibrated at 45% B (37–40 min). ESI source parameters were as follows. In positive mode (ESI+): sheath gas 15 (arb. units), spray voltage 4.5 kV, capillary temperature 330 °C, capillary voltage 48 V, and tube lens 105 V. In negative mode (ESI−): sheath gas 15 (arb. units), spray voltage 5.5 kV, capillary temperature 330 °C, capillary voltage −47 V, and tube lens −85.6 V. Mass spectra were acquired in full-scan mode over m/z 110–1800. Negative mode was used for PG and CL detection, while positive mode supported the detection of lysylated species and glycolipids. Data-dependent MS/MS experiments were performed using a normalized collision energy of 30% and an isolation width of 3 m/z to obtain diagnostic fragment ions for lipid identification. This LC–MS/MS approach enabled the detection of membrane lipid remodeling, particularly the incorporation of lysyl groups into phosphatidylglycerols and variations in glycolipid composition under the studied conditions.

### 2.7. Carotenoids analysis by LC-DAD-APCI-MS/MS

Carotenoids were identified and quantified using a previously reported method with minor modifications. Dried extracts were reconstituted at 10,000 ppm in MeOH:CHCl3 (2:1, v/v) and analyzed using an UHPLC Dionex UltiMate 3000 system equipped with a diode-array detector (DAD) coupled to an LCQ Fleet ion trap mass spectrometer (Thermo Scientific) via an atmospheric pressure chemical ionization (APCI) source operated in negative mode. Data acquisition and processing were performed using Xcalibur 4.3 software. The chromatographic separation was carried out on a YMC-C30 column (150 × 4.6 mm, 3 μm; YMC America) with a C18 guard cartridge (4 × 2 mm, 3 μm), at room temperature. A 10 μL injection volume was used, with samples maintained at 5 °C in the autosampler. The mobile phases consisted of (A) methanol:methyl tert-butyl ether:water (80:18:2, v/v/v) and (B) methanol:methyl tert-butyl ether:water (8:89:3, v/v/v), both containing 400 mg/L ammonium acetate. The flow rate was 450 μL min⁻¹.

The gradient elution was as follows: 5% B (0–3 min), 5–10% B (3–9 min), 10–25% B (9–19 min), 25–40% B (19–23 min), held at 40% B (23–27 min), increased to 100% B (27–31 min), held at 100% B (31–34 min), returned to 5% B (34–36 min), and equilibrated at 5% B until 40 min. DAD data were acquired over the UV–Vis range (240–600 nm), with chromatograms extracted at 465 nm for carotenoids and 269 nm for menaquinones (MK). The APCI source parameters were: vaporizer temperature 300 °C, discharge current 15 μA, capillary voltage −25 V, tube lens −80 V, capillary temperature 350 °C, sheath gas 50 (arb. units), and auxiliary gas 30 (arb. units). Mass spectra were acquired in full-scan mode (m/z 65–1200), with data-dependent MS/MS performed at 30% collision energy and an isolation width of 3 m/z to generate characteristic fragment ions. Carotenoids were putatively identified based on UV–Vis absorption spectra and APCI-MS/MS fragmentation patterns, in agreement with previously reported data.

For quantification, β-carotene was used as an external standard due to the lack of commercially available standards for *S. aureus* carotenoids. Calibration curves were constructed over the range 1–10 μg mL⁻¹ (y = 175737x − 101826; R² = 0.999). A molecular weight correction factor was applied to account for differences in detector response, following previous reports. Carotenoid concentrations were expressed as μg g⁻¹ dry cell weight.

### 2.8. Fourier transform infrared (FTIR) spectroscopy

The experiments were prepared on a BioATR II cell integrated to a Tensor II spectrometer (Bruker Optics, Ettlingen, Germany) with a liquid nitrogen MCT detector using a spectral resolution better than 0.4 cm^-1^ and 120 scans per spectrum. The temperature range was set by a computer-controlled circulating water bath Huber Ministat 125 (Huber, Offenburg, Germany). All the experiments were carried out with a heating rate of 1 °C/min and a stabilization time of 120 seconds at each temperature. First, the background was taken using buffer (HEPES 20 mM, 500mM NaCl and 1mM EDTA) from 10 to 45 °C. Subsequently, the lipid extracts were deposited on the silicon crystal (0.3 mg) to prepare *in situ* on the BioATR II cell the solid-supported lipid bilayers, hydrated with 20 µL of buffer at 37 °C for 10 min, and measurements were recorded at each temperature of the ramp. Finally, to determine the position of the vibrational band in the range of the second derivative of the spectra, all the absorbance spectra were cut in the 2970 - 2820 cm^-1^ range, shifted to a zero baseline and the peak picking function included in OPUS 8.8.4 software. The results were plotted as a function of the temperature using the OriginPro 8.0 software (OriginLab Corporation, USA). To determine the melting temperature Tm of the lipids, the curve was fitted according to the Boltzmann model to calculate the inflection point of the obtained thermal transition curves using Python.

## 3. Results

### 3.1. The structure of LL-37 and ATRA-1 antimicrobial peptides

Fig. 1. shows the sequences of LL-37 and ATRA-1. Helical wheel projections show the amphipathic distribution of residues for both peptides. The human cathelicidin contains a precursor domain followed by LL-37 in its C-terminus. Proteolytic cleavage of the precursor domain leads to an active LL-37 [47], LL-37 is in random coil conformation in pure aqueous solution, and assumes an amphipathic alpha helical configuration in the presence of lipid bilayers with a disordered C-terminal tail that does not form an amphipathic α-helix [48, 49]. LL-37 binds to both anionic and zwitterionic lipid bilayers, binding parallel to the surface at low concentrations [50, 51] and presenting a higher propensity for insertion into a vertically inserted conformation in anionic bilayers [51]. LL-37 forms tetrameric toroidal pores that lead to bacterial membrane disruption [17, 18]. In addition to this membrane disruption mechanism, LL-37 has been shown to regulate the immune response and is widely present in diverse tissues in the human body. In particular, LL-37 is found in epithelial cells of the human lung, and has broad antimicrobial activity in the airway surface, where *S. aureus* can reside [52]. LL-37 is also produced by neutrophils that are recruited to the mucosa [53], another site prone to *S. aureus* colonization [54]. It is evident that *S. aureus* is normally exposed to LL-37 in human tissue, raising the question of how the membrane of this pathogen may adapt to the presence of the human cathelicidin.

**Fig. 1.**
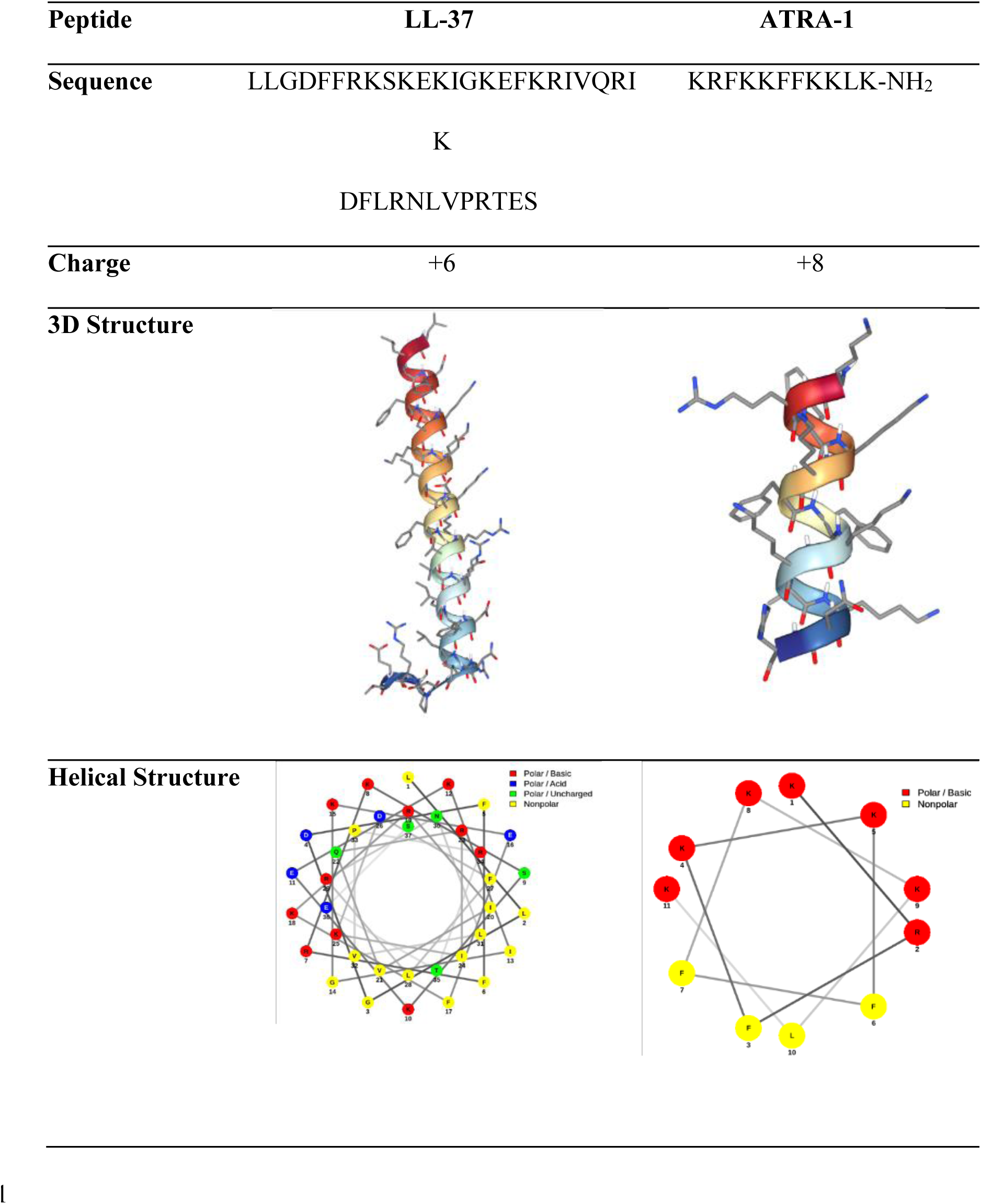
Amino acid sequences, net electric charge, 3D structures (generated with PEP-FOLD 4), and helical wheel projections (generated with PEP-FOLD 3) of the peptides LL-37 and ATRA-1, used to adapt *S. aureus* to antimicrobial peptide exposure.

ATRA-1 is an antimicrobial peptide identified within the sequence of the Chinese cobra (*Naja atra*) cathelicidin. The peptide assumes a very well-defined α-helical amphipathic structure when exposed to SDS, with an exclusively cationic and hydrophobic face [55] as shown in the wheel structure representation (Fig. 1). ATRA-1 shows strong antimicrobial activity, including against *S. aureus*, without presenting hemolytic activity [55, 56]. ATRA-1 is a short peptide with a size of 11 residues, resulting in an α-helical length of approximately 16.5 Å, which is less than half of the expected width of the bilayer membrane [23]. This implies that ATRA-1, in contrast to LL-37, does not completely traverse the membrane when inserted vertically. Instead of disrupting the bacterial membrane through the formation of a multimeric pore, the mechanism of membrane disruption is more in line with a carpet-like mode of action.

### 3.2 Electron microscopy images of S. aureus in the presence of sub-lethal concentrations of LL-37 and ATRA-1

To corroborate the antimicrobial activity of LL-37 and ATRA-1 in *S. aureus,* cells were examined via scanning electron microscopy (SEM) at two time points: 40 minutes and 24 hours post-exposure. Due to sample volume requirements and peptide availability, experimental conditions had to be adjusted for each time point. Short-term exposure (40 min) utilized a higher inoculum density, 10 μL of bacterial saturated culture in 1 mL of media, and elevated peptide concentrations (LL-37: 400 μg/mL; ATRA-1: 300 μg/mL). Conversely, long-term samples (24 h) followed standard protocols of 1 μL of culture in 10 ml of media, with lower concentrations (LL-37: 40 μg/mL; ATRA-1: 30 μg/mL). All samples were grown under stirring conditions (150 rpm) at 37 °C. After both 40 minutes and 24 hours after exposure, cells were fixed and observed through SEM.

When *S. aureus* is treated with the elevated concentrations of LL-37 and ATRA-1, extensive damage is observed in the integrity of the bacterial membrane, as shown in Fig. 2. The damage appears in a large proportion of bacteria and is observed as cells exhibiting localized membrane collapse and irregular cell morphologies. Both peptides induce qualitatively similar levels of membrane damage. This shows only that both peptides are active in *S. aureus*. SEM images of *S. aureus* fixed after 24 hours of growth in the presence of sub-lethal concentrations of LL-37 and ATRA-1 showed round cells, and dividing cells with no evidence of membrane damage, with similar morphologies as untreated bacteria (Fig. 2). *S. aureus* has many defense mechanisms against antimicrobial peptide activity that include the generation of exopolymers [57–59] and the secretion of extracellular proteases [58, 59]. However, in this study we wanted to focus on changes in lipid composition in response to antimicrobial peptide activity.

**Fig. 2.**
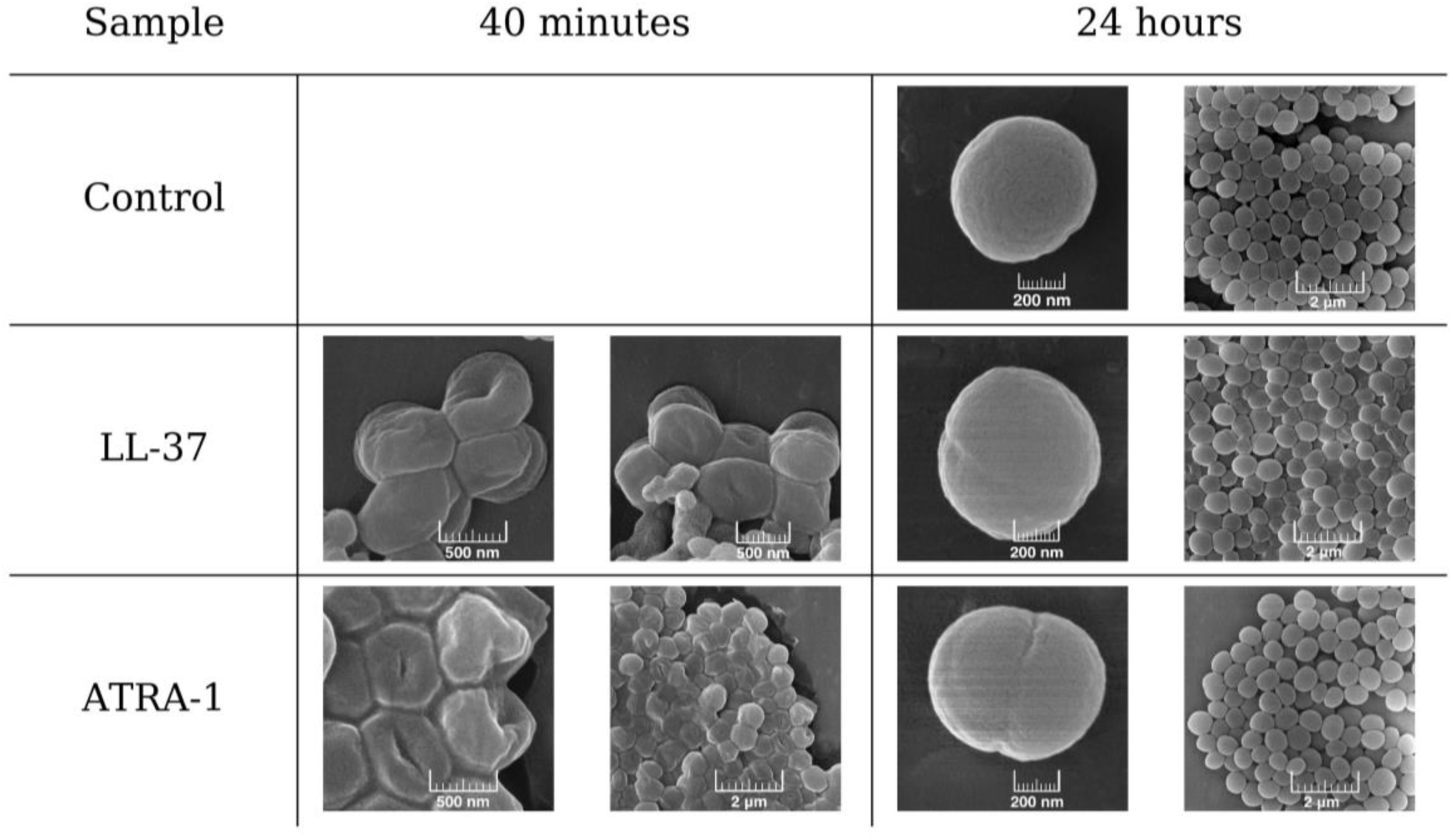
Scanning Electron Microscopy (SEM) images showing the morphological alterations in *S. aureus* cells 40 minutes after exposure to LL-37 and ATRA-1, and subsequent adaptation 24 hours after exposure. Untreated control cells are shown for comparison.

### 3.3 Changes in phospholipid and glycolipids composition and membrane surface potential of S. aureus membranes in the presence of sub-lethal concentrations of LL-37 and ATRA-1

Phospholipid composition in *S. aureus* treated with LL-37 and ATRA-1 was analyzed by liquid chromatography–electrospray ionization tandem mass spectrometry (LC-ESI-MS/MS) for all growth conditions. Figure 3 shows the general chemical structure (Fig. 3a) and the variations in the proportion (Fig. 3b) of each of the main phospholipid and glycolipid components in *S. aureus*: PG, CL, Lysyl-PG, MGDG, and DGDG when exposed to LL-37 and ATRA-1.

**Fig. 3.**
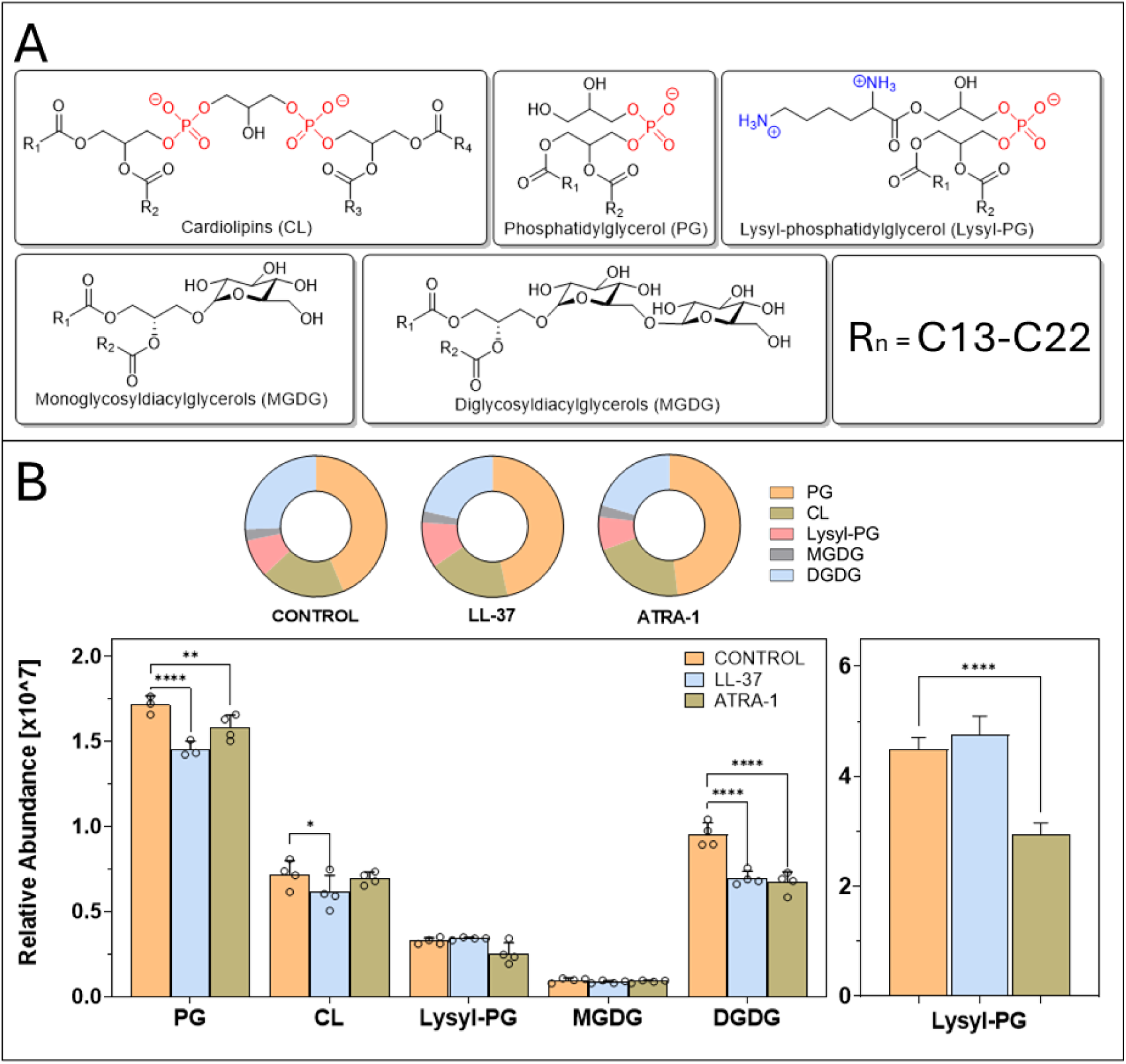
(a) Molecular structures of the five major membrane lipid components of *S. aureus*: the phospholipids phosphatidylglycerol (PG), lysyl-phosphatidylglycerol (Lysyl-PG), and cardiolipin (CL), and the glycolipids monoglucosyldiacylglycerol (MGDG) and diglucosyldiacylglycerol (DGDG). (b) Relative lipid composition of control and peptide-treated samples. Donut charts show the proportional distribution of PG, CL, Lysyl-PG, MGDG, and DGDG. Bar plots show the relative abundance of each lipid class detected in negative ion mode (ESI−). Lysyl-PG relative abundance obtained in positive ion mode (ESI+) is shown separately on the right. Data are presented as mean ± SD (n = 4). Statistical significance was determined using two-way ANOVA followed by Dunnett’s multiple comparison test. Significance levels: *P < 0.05, **P < 0.01, ***P < 0.001, \*\***P < 0.0001.** Outliers were identified and removed using the Grubbs test (2 values).

The main phospholipid component found in *S. aureus* membranes is the anionic phospholipid PG, although it can vary widely with the strain and growth conditions of the bacteria [60]. Cardiolipin is synthesized by attaching phosphatidic acid to a PG molecule through the glycerol headgroup [24]. This is done through an enzymatic process mediated by Cls1 and Cls2 [27]. CL levels can vary in response to osmotic stress and high salinity [61] and changes in oxygen levels [64], maintaining levels of approximately 20 mol% of phospholipid content under standard growth conditions [62]. Cardiolipin increases membrane rigidity and has been shown to protect the membrane from the pore-forming activity of antimicrobial peptides [21]. The cationic phospholipid Lysyl-PG regulates the membrane surface potential and has been associated with increased resistance to antimicrobial peptides through a reduction in membrane anionic charge that leads to lower binding affinities of antimicrobial peptides [62, 63].

In addition to PG and CL, DGDG and MGDG are the most prevalent lipid species present in *S. aureus* membranes [39], [40], and incorporation of oleic acid into these lipids, which promotes conicity in MGDG and DGDG, leading to the emergence of a melting event characteristic of a non-lamellar phase transition [39].

Treatment with LL-37 and ATRA-1 does not lead to statistically significant increases in CL with respect to PG for either peptide treatment. Even though CL has been shown to increase resistance to antimicrobial peptide activity [21], the results in Fig. 3b suggest that no regulation of CL levels occurs in the presence of these two antimicrobial peptides.

It is worth noting that membranes of *S. aureus* present basal levels of Lysyl-PG. Therefore, *S. aureus* membranes are not fully anionic under standard growth conditions for wild-type strains. Basal levels of Lysyl-PG have been reported previously [62]. This point is of relevance since models for Gram-positive bacteria used in the evaluation of antimicrobial peptides emphasize the use of 100 mol% anionic lipids. Fig. 3b shows that Lysyl-PG levels are sensitive to treatment with ATRA-1 but not LL-37. Treatment with LL-37 induces no significant changes in Lysyl-PG levels, while ATRA-1 induces a decrease in Lysyl-PG levels. This emphasizes that Lysyl-PG levels are not simply positively correlated to the presence of antimicrobial peptides, and that other biophysical aspects in the bilayer membrane play a role.

The presence of LL-37 and ATRA-1 leads to a reduction in DGDG levels in the membrane which suggests an increase in negative curvature stress related to an increase in the MGDG/DGDG ratio. This result points to the possible correlation between exposure to antimicrobial peptides and changes in negative curvature stress in the membrane [39]. Increases in negative curvature stress have been associated with increased resistance to antimicrobial peptide activity [39].

### 3.4 Phospholipid acyl chain length distributions remain constant in the presence of sublethal levels of LL-37 and ATRA-1

Fig. 4 shows the acyl chain length distributions for PG, CL and Lysyl-PG phospholipid species for control, and after treatment with sublethal concentrations of LL-37 and ATRA-1. The lipid profile of *S. aureus* is characterized by a predominance of odd-numbered, branched-chain fatty acids, consistent with previous reports. The most abundant species detected were consistent with branched anteiso fatty acid compositions (e.g., anteiso-C15:0/C17:0; total carbon number 32:0) for PG and Lysyl-PG, while CL species were predominantly composed of combinations consistent with four acyl chains totaling 64:0. The presence of sublethal concentrations of LL-37 and ATRA-1 does not induce significant changes in the three lipid species. Only CL has a slight decrease in the short-chained species complemented with a slight increase in the 66:0 species. In general, acyl chain composition is not sensitive to the treatment with these antimicrobial peptides.

**Fig. 4.**
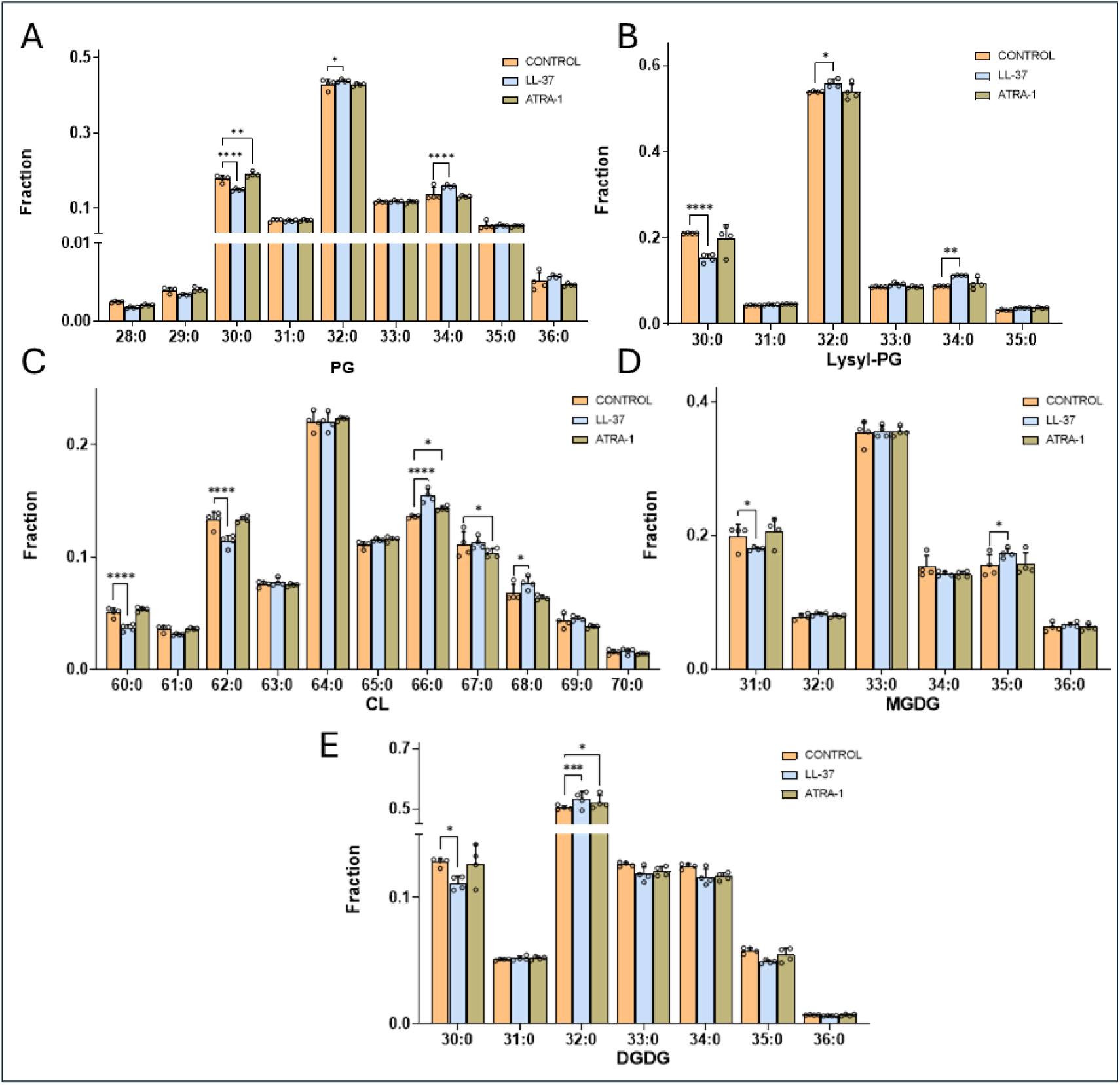
Distribution of acyl chain lengths of the five major membrane lipids of *S. aureus*: the phospholipids phosphatidylglycerol (PG), lysyl-phosphatidylglycerol (Lysyl-PG), and cardiolipin (CL), and the glycolipids monoglucosyldiacylglycerol (MGDG) and diglucosyldiacylglycerol (DGDG) for untreated, LL-37 treated, and ATRA-1 treated *S. aureus* cells.

### 3.5 Changes in carotenoid and menaquinone composition in S. aureus in the presence of sublethal concentrations of LL-37 and ATRA-1

Changes in carotenoid content and composition in the presence of sublethal concentrations of LL-37 and ATRA-1 were analyzed in Fig 5. Carotenoids are lipid pigments that provide *S. aureus* with its characteristic golden color. These molecules protect *S. aureus* against oxidative stress due to their ability to act as free-radical scavengers. Carotenoid levels are controlled through the levels of oxygen present in the growth media, through the two-component system AirRS [65]. Carotenoids also provide mechanical resistance [32] and have been shown to protect *S. aureus* against antimicrobial peptide activity [32, 66]. The synthesis of carotenoids is regulated by the *crtOPQMN* operon, where the end-product of the metabolic pathway is staphyloxanthin (STX) [31, 33], which accumulates in the membrane in the stationary phase [32]. Figure 5a shows the general chemical structure of the STX molecule. The molecule is composed of a glucose headgroup with two side chains. The first is a triterpenoid side chain that presents a series of conjugated double bonds, which leads to a rigid straight chain. The second side chain is a saturated acyl chain with varying lengths. The enzyme CrtQ attaches a glucose molecule to the triterpenoid 4,4’- diaponeurosporenoic acid (4,4’-DNPA), followed by the attachment of an acyl chain through CrtO. It has been shown that 4,4’-DNPA and STX are in chemical equilibrium in *S. aureus* in the stationary phase [33]. Figure 5b shows the relative abundance of several putatively identified STX-related species (STX-Iso-4, STX-Iso-5, STX-Iso-6, and STX C17 Iso-1), as well as the precursor 4,4′-DNPA. Differences in carotenoid content associated with pigmentation were qualitatively assessed during the lipid extraction. The test tubes are shown in Figure 5a inset, where the LL-37 exposed bacteria have a clearly paler orange tone with respect to control, and the ATRA-1 exposed bacteria only show slight differences in the color of the bacterial population with respect to control.

**Fig. 5.**
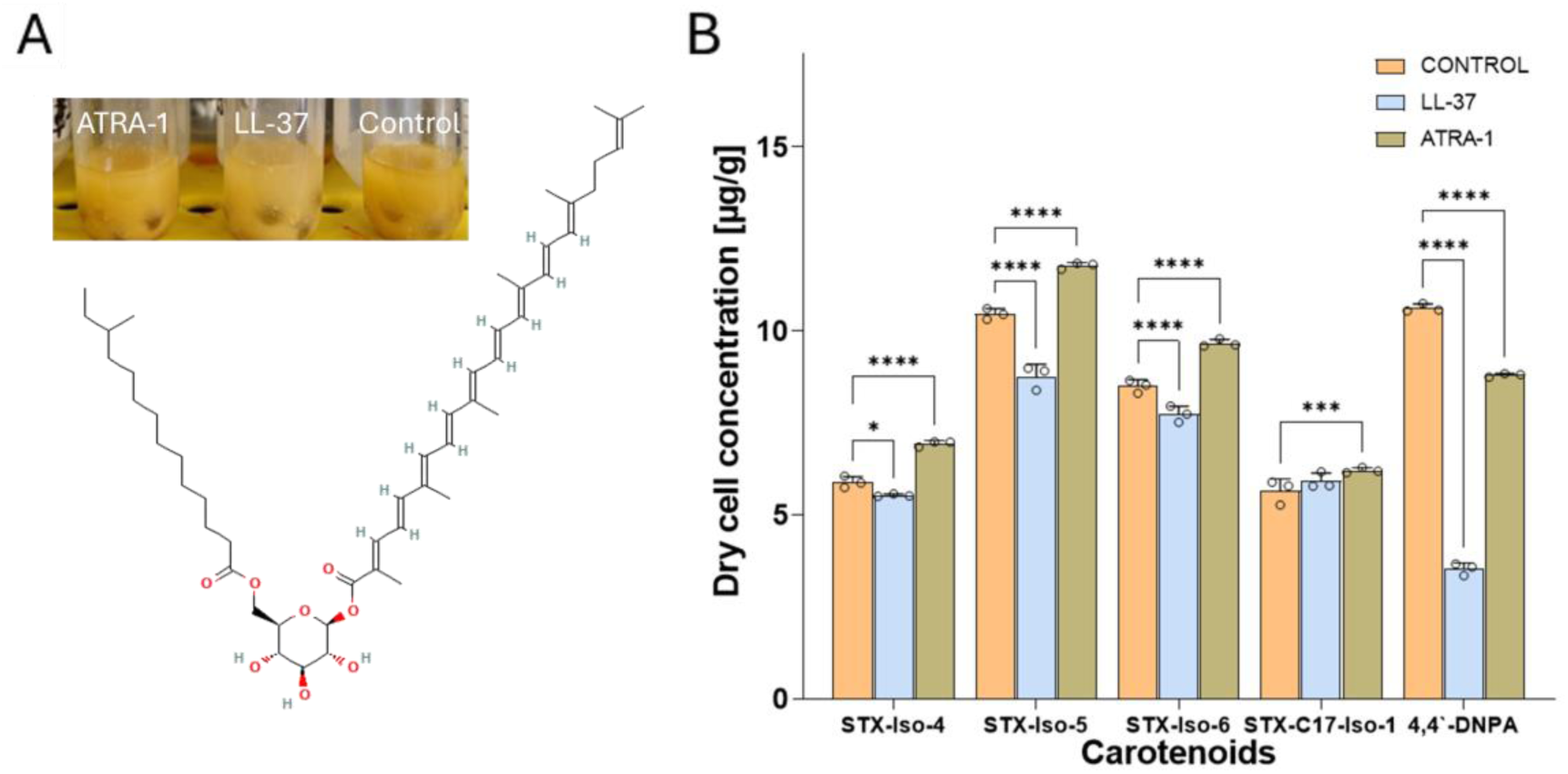
(a) Structure of the *S. aureus* carotenoid staphyloxanthin. Inset: bacterial pellets show visibly paler coloration for LL-37-exposed cells. (b) Dry weight concentrations of three STX-C15, a STX-C17 isoforms and the 4,4’-DNPA precursor for untreated, LL-37 treated, and ATRA-1 treated *S. aureus* cells. Data are presented as mean ± SD (n = 3). Statistical significance was determined using two-way ANOVA followed by Dunnett’s multiple comparison test. Significance levels: *P < 0.05, **P < 0.01, ***P < 0.001, \*\***P < 0.0001.**

Measurement of carotenoid content (Fig. 5b) shows a reduction of several carotenoid species in the LL-37 treated samples with respect to control, including all STX species, except for STX C17 Iso-1. In addition, a sharp decrease in the precursor 4,4’-DNPA, indicates an overall reduction in carotenoid production when *S. aureus* is treated with sublethal concentrations of LL-37. The trend is not as clear in the case of ATRA-1 exposed bacteria which presents an overall increase for all STX species except for STX C17 Iso-1, and a slight decrease in the 4,4’-DNPA precursor content with respect to control. For LL-37 treated samples it is clear that staphyloxanthin production is being downregulated.

Menaquinones (MKs) are isoprenoid quinones (Fig. 6a) located in the cytoplasmic membrane of *S. aureus* that participate in the oxidative phosphorylation pathway leading to the production of ATP. Menaquinones take part in the electron transport chain by transferring electrons derived from NADH oxidation to the heme groups found in cytochromes. Under anaerobic conditions, menaquinones participate in non-oxidative metabolic pathways involving nitrogen. Targeting menaquinone synthesis has been proposed as a strategy for reducing *S. aureus* virulence. However, defects in menaquinone synthesis have been identified in small colony variants (SCV) of *S. aureus*, which have been associated with persistent infections [41]. We report three menaquinones (MK7, MK8, and MK9), where the most prevalent MK species in *S. aureus* is MK8, consistent with previous reports [67]. Both antimicrobial peptides induce a reduction in all MK species, with a more accentuated change in the case of LL-37 treatment (Fig. 6b).

**Fig. 6.**
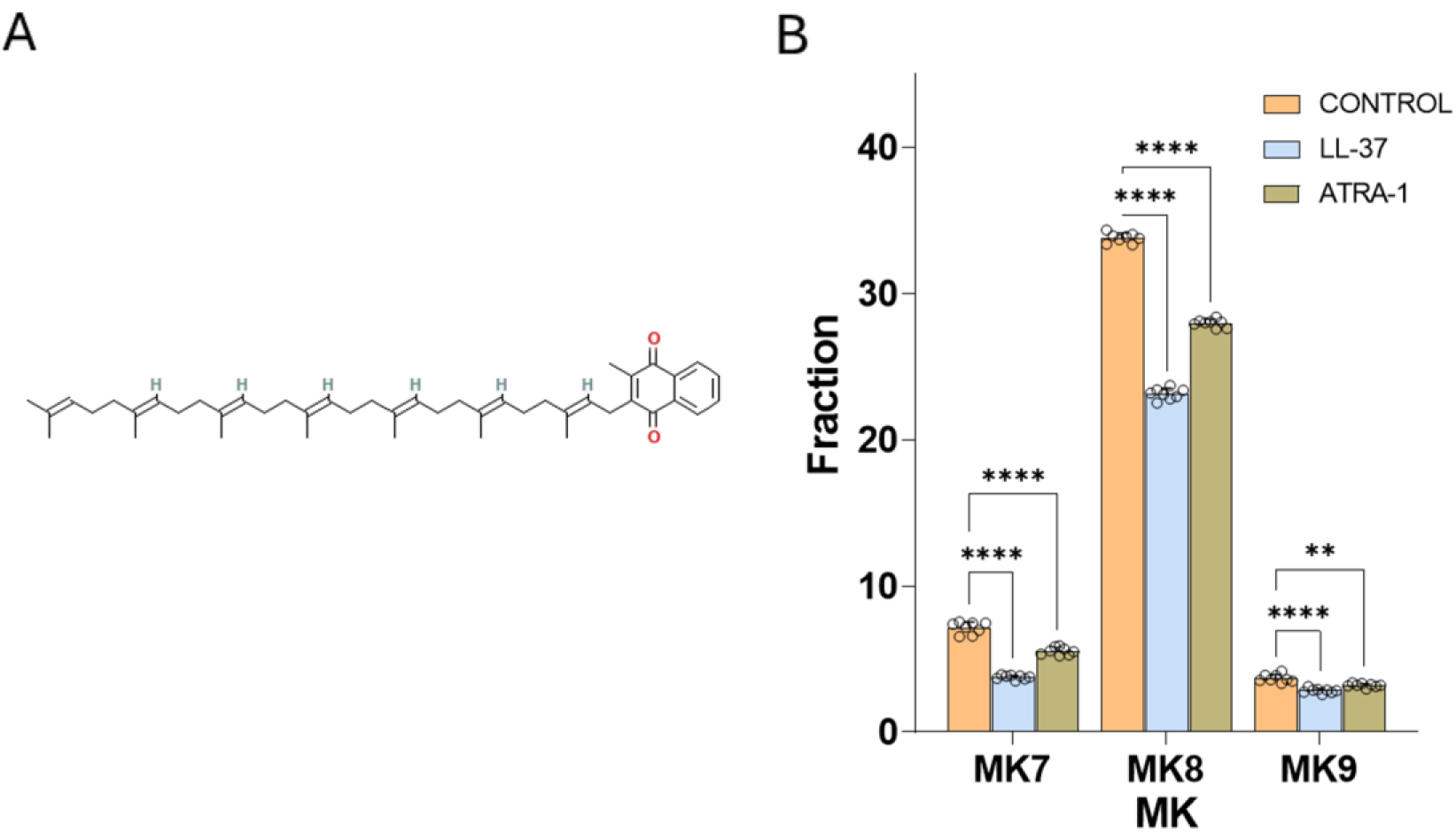
(a) Chemical structure of menaquinone-7 (MK-7). (b) Integrated peak areas of MK-7, MK-8, and MK-9 for untreated, LL-37 treated, and ATRA—1 treated *S. aureus* cells. Data are presented as mean ± SD (n = 8). Statistical significance was determined using two-way ANOVA followed by Dunnett’s multiple comparison test. *P < 0.05, **P < 0.01, ***P < 0.001, ****P < **0.0001.**

### 3.6 Surface potential measurements of total lipid extracts treated with sublethal concentrations of LL-37 and ATRA-1

Z-potential measurements were conducted on total lipid extracts from *S. aureus* control, LL-37 exposed, and ATRA-1 exposed cells (Fig. 7). An estimate of the surface charge density for the different treatments was calculated using Equation 1 and presented in Table 1. Lipid extracts were extruded to form unilamellar vesicles with sharp size distributions centered between 100 and 150 nm. Zeta potential measurements showed a significant drop in magnitude of the negative surface potential in the LL-37 exposed bacteria, and a significant increase in magnitude of the negative surface potential in the ATRA-1 exposed bacteria, with respect to control (Fig. 7). The changes are in line with the observed changes in Lysyl-PG levels. However, the decrease in the magnitude of the negative surface potential in the case of LL-37 appears more significant than what would be attributed to the change in Lysyl-PG levels, which do not vary significantly with LL-37 treatment. Other lipid components, including neutral species, may be playing a role in modulating the surface potential in the presence of LL-37. In particular, the observed drop in diaponeurosporenoic acid levels associated with staphyloxanthin synthesis (Fig. 5) is consistent with the decrease in anionic surface potential magnitude observed for LL-37-treated samples. The diversity of lipid species present in *S. aureus* (Fig. 3, 5 and 6) indicates that the lipid bilayer surface charge can be modulated through changes in various lipid components. A calculation of the surface charge density is presented using the Grahame equation that takes into account ionic screening effects.

**Fig. 7.**
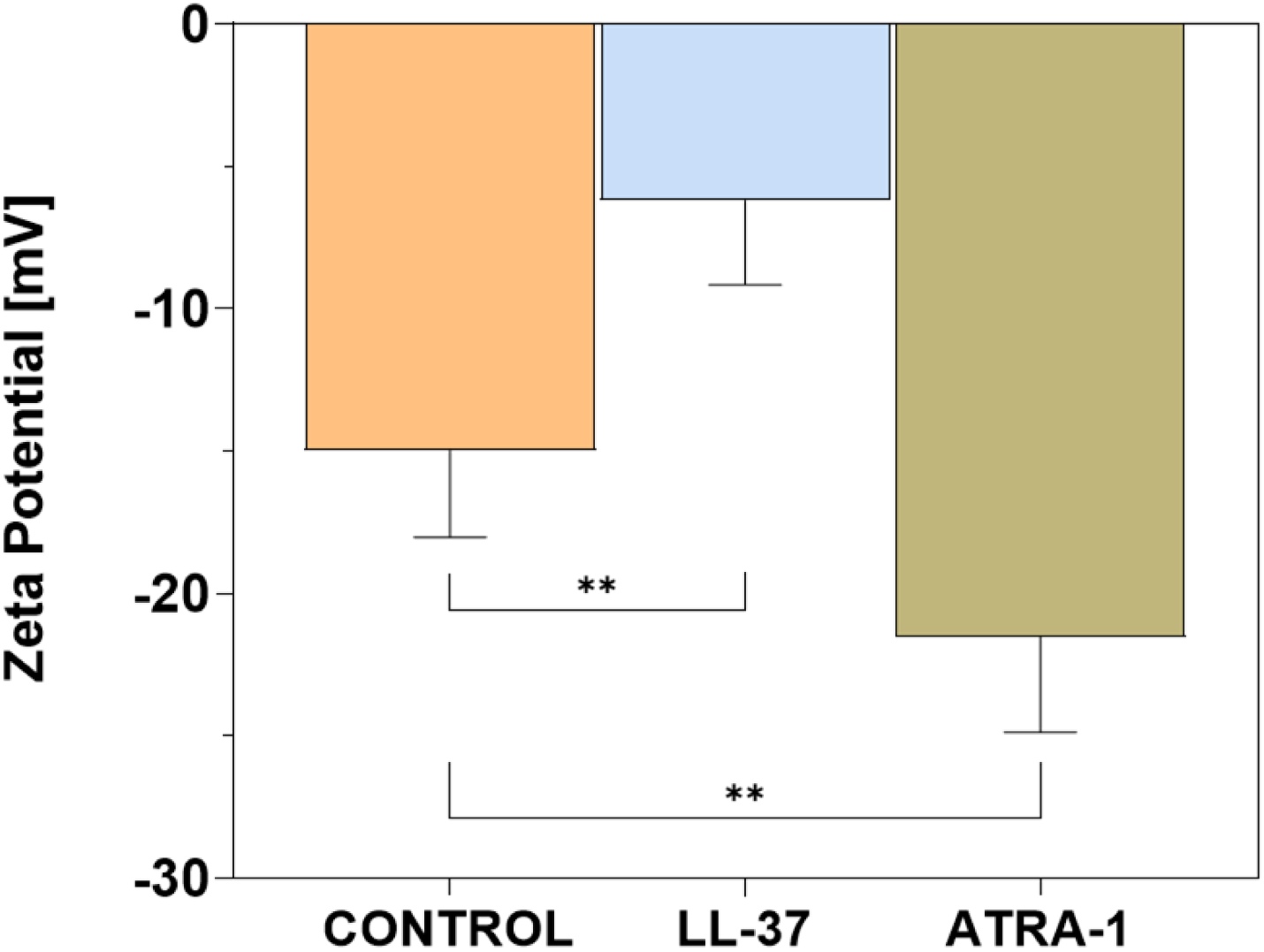
Changes in the membrane surface potential of 100 nm unilamellar vesicles composed of total lipid extracts for untreated, LL-37 treated, and ATRA-1-treated *S. aureus* cells.

**Table 1.** Calculation of surface charge density based on the measured Zeta Potential for total lipid extracts from untreated, LL-37 treated, and ATRA—1 treated *S. aureus* cells.

| Group | Zeta Potential ($\zeta$)<br>[mV] | Surface Charge Density ($\sigma$)<br>[C/m<sup>2</sup>] |
| --- | --- | --- |
| Control | -14.98 ± 3.04 | -0.0152 ± 0.0032 |
| LL-37 treated | -6.21 ± 2.96 | -0.0062 ± 0.0030 |
| ATRA-1 treated | -21.52 ± 3.36 | -0.0221 ± 0.0037 |

### 3.7 Gel to liquid-crystalline phase transitions in S. aureus total lipid extracts in the presence of sublethal concentrations of LL-37 and ATRA-1

To study the thermodynamic lipid phase behavior of the membrane, the υsCH2 acyl chain vibration was monitored as a function of temperature by FTIR. *S. aureus* membranes present gel (Lβ) to liquid-crystalline (Lα) cooperative phase transitions, centered around 15°C [32, 33, 44, 65]. The phase transition temperature has been shown to be sensitive to growth conditions [64]. A key regulator of this phase behavior is staphyloxanthin [38], where increases in STX induce a decrease in the phase transition temperature due to the destabilization of the gel phase, and an increase in the level of lipid packing in the liquid-crystalline phase, as reflected by an increase in the υsCH2 vibration wavenumber. An inverted hexagonal phase transition has been reported recently above the growth temperature as measured by differential scanning calorimetry [39].

The thermotropic behavior of the υsCH2 vibration for untreated, LL-37 treated, and ATRA-1 treated cells is plotted in Figure 8a. Here, the increase of this number denotes a higher acyl-chain disorder amongst the lipids, and the inflection points of the curves mark the transition temperature between the gel phase and the liquid-crystalline phase, denoted as melting temperature (Tm) [38]. To measure the phase transition temperature, the first derivative of the wave number with respect to the temperature is plotted (shown in Fig. 8b), where the location of the maximum in the first derivative denotes the Tm.

**Fig. 8.**
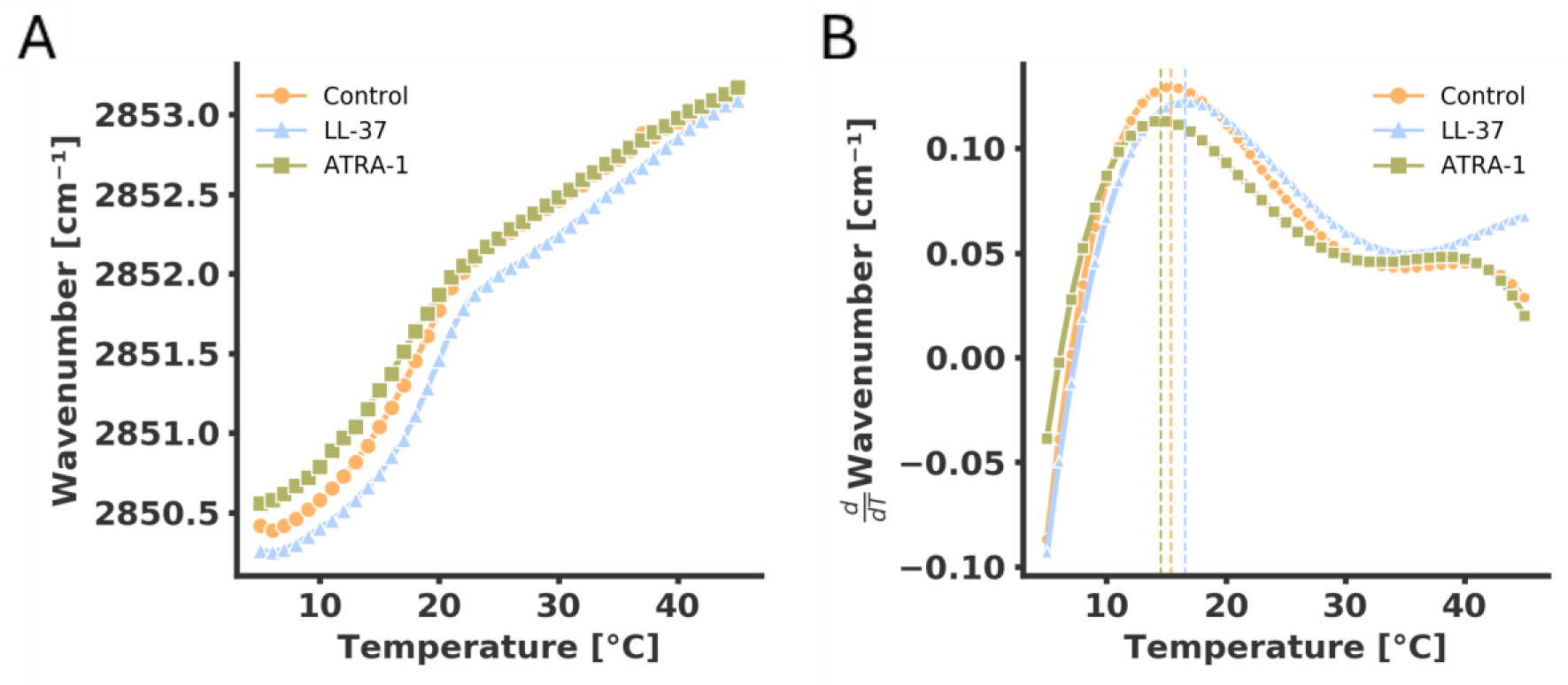
(a) Thermotropic plots of the νCH2 symmetric stretching vibrations band of the acyl chain methylene groups as measured by FTIR showing changes in the lipid phase behavior of *S. aureus* total lipid extracts. (b) First derivative of wavenumber vs. temperature curve, highlighting the melting transitions (Tₘ) for untreated (15.38 °C), LL-37 treated (16.58 °C) and ATRA-1 treated (14.51 °C) cells.

The LL-37 treated sample shows an increase in the phase transition temperature coupled by a decrease in the υsCH2. This is consistent with a reduction in the levels of carotenoids in the LL-37 treated samples (Fig. 5b), visually apparent as shown in the reduction in coloration levels. The same trend has been shown to occur in STX biosynthesis knock-outs and in synthetic systems [33, 38]. This implies that the liquid-crystalline phase becomes less rigid for the LL-37 treated cells at physiological temperatures (37 °C). In the case of ATRA-1, total lipids show a slight decrease in the phase transition temperature (Fig. 8b).

## 4. Discussion

This study surveys several lipid components of the membrane of *Staphylococcus aureus* in response to exposure to two antimicrobial peptides, LL-37 and ATRA-1. The two peptides have different origins. LL-37 is a human cathelicidin, and *S. aureus* is exposed to the human cathelicidin in its ecological niche, such as the mucosa and skin of the human body. LL-37 is long enough to traverse the membrane, forming tetrameric pores. In contrast, ATRA-1 is a short peptide derived from snake venom that does not traverse the membrane but acts in a carpet-like membrane disruption mechanism. Clear differences are observed with regards to changes in lipid composition of *S. aureus* membranes when treated with LL-37 and ATRA-1, indicating peptide specific differences in the physiological response during adaptation of the bacterial membrane. There are three main findings that we would like to emphasize in this discussion: (1) the downregulation of carotenoid production in the presence of LL-37, (2) reduced production of menaquinones induced by both peptides, and (3) a change in the MGDG/DGDG ratio induced by both antimicrobial peptides.

The presence of LL-37 induces a clear reduction in carotenoid content (Fig. 5), where the greatest effect is observed in the precursor 4,4’-DNPA, which is significantly reduced. ATRA-1 does not induce this decrease. Staphyloxanthin and 4,4’-DNPA, have been shown to be in equilibrium in the stationary phase [32], so a reduction in 4,4’-DNPA is an indication of an overall reduction in carotenoids. With regards to the biophysical properties of the membrane, carotenoids increase the rigidity of the membrane in the liquid-crystalline phase [32, 38, 64]. A reduction in carotenoid content induced by LL-37 is reflected in a change in the gel to liquid-crystalline phase transition temperature (Fig. 8), which has been reported to occur when carotenoid content varies [33, 38]. This is also coupled with a change in pigmentation levels observed in the LL-37 treated samples (Fig. 5a). In general, LL-37 affects staphyloxanthin levels in *S. aureus*, influencing the biophysical properties of the bacterial membrane.

Inhibition of carotenoid synthesis has been proposed as a strategy for increasing *S. aureus* susceptibility to antibiotics or free radicals generated through an immune response [34, 37, 69]. The presence of staphyloxanthin not only provides mechanical resistance to antimicrobial peptide activity [32, 64], but also protects the bacterial membrane from free radicals generated by macrophages and neutrophils. A reduction in carotenoids is likely to result in increased susceptibility of *S. aureus* to the immune response. It is therefore notable to observe that LL-37 reduces carotenoid levels in S*. aureus.* It has been reported that LL-37 acts as an immunoregulatory agent that can increase ROS production and promote engulfment of S. aureus in neutrophils [70] and regulate chemokine production in a selective manner [70, 71]. In general, LL-37 acts as a multi-factor regulator of the immune response in addition to its role in generating pores in bacterial membranes.

Many regulators participate in modulating the production of staphyloxanthin in *S. aureus*. For example, the global stress response regulator SigB recruits RNA polymerases to the promoter that drives the transcription of the *crtOPQMN* operon required for synthesis of staphyloxanthin [36, 72]. SigB is activated through two-component systems such as SaeRS [73]. Other two component systems such as MsaB [74] and YjbIH [75] upregulate staphyloxanthin production and are involved in bacterial virulence and survival against host immune cells [73]. In particular, AirSR is a two-component system that detects oxygen levels through a heme binding domain and acts in an independent manner to SigB to upregulate staphyloxanthin production to increase tolerance to oxidative stress [65]. The downregulation of staphyloxanthin and its precursor 4,4’-DNPA may involve the disruption of one or more components of this regulatory network through the interaction of LL-37 with the bacterial membrane. Since staphyloxanthin production is an important virulence factor in *S. aureus,* due to its protective role during oxidative stress and antimicrobial peptide activity, this downregulation induced by the human cathelicidin LL-37 is worth exploring in future work.

Both ATRA-1 and LL-37 induce a reduction in menaquinone levels (Fig. 6). *S. aureus* uses menaquinones (MKs) to transfer electrons during oxidative phosphorylation, where a reduction in menaquinone levels normally leads to reduced metabolic rates due to deficient respiration. Reduced menaquinone biosynthesis has been shown to lead to the small colony variant (SCV) phenotype in *S. aureus.* Typically, electron-transport deficient SCVs can be restored to normal colony growth by supplementing with MK [76]. MK deficiencies are therefore related to reduced growth rates. Electron-transport deficient SCV have been isolated from patients with chronic infections typically presenting methicillin resistance, and SCVs have been shown to present increased resistance to antibiotic treatment [77, 78]. SCV can also be induced through environmental pressure such as treatment with gentamicin [79, 80]. Reduced production of menaquinones is therefore relevant in the understanding of how antimicrobial peptides such as LL-37 and ATRA-1 interact with *Staphylococcus aureus,* and whether antimicrobial peptides can trigger a low growth state with reduced menaquinone production.

The glycolipids MGDG and DGDG are in high proportion in *S. aureus* membranes (Fig. 3b), playing a key role in regulating biophysical properties. In a recent publication, their role in modulating the mechanical curvature stress in *S. aureus* membranes was highlighted [39]. Both lipids, but more prevalently MGDG, leads to increased negative curvature stress in the membrane due to their inverted conical shape. This curvature stress can play an important role in cell division since lipids possessing negative curvature can reduce the free energy requirement for forming the division septum. Additionally, negative curvature stress has been shown to inhibit the pore forming activity of antimicrobial peptides [39]. It is worth noting that CL also plays a similar role in increasing negative curvature in the membrane due to its conical shape. Fig. 3b shows that the largest proportion of glycolipids corresponds to DGDG, and that this lipid is reduced significantly in the presence of LL-37 and ATRA-1. Since the fractions of MGDG and CL do not change significantly with antimicrobial peptide treatment, the reduction in DGDG would suggest an increase in negative curvature stress in the membrane.

## 5. Conclusions

A lipidome response is observed in *S. aureus* membranes when exposed to the antimicrobial peptides LL-37 and ATRA-1. However, the changes in lipid composition are dependent on the type of peptide. Overall, the human cathelicidin LL-37 induces a decrease in carotenoid production, while the snake derived antimicrobial peptide ATRA-1 induces a decrease in the cationic phospholipid Lysyl-PG. The decrease in carotenoids induced by LL-37 stands out in the context of carotenoids acting as virulence factor, where carotenoids provide *S. aureus* with a defense mechanism towards free radicals generated in the immune response, and increased membrane rigidity to counteract antimicrobial peptide activity. It is worthwhile investigating whether LL-37 can downregulate carotenoid production in *S. aureus* to make it more susceptible to the immune response. Both peptides induce a reduction in menaquinones, which take part in electron transfer during oxidative phosphorylation. Since deficiencies in menaquinone levels lead to small colony variants that have been shown to be persistent to antibiotic treatment, further research is needed to investigate if small colony variants can be induced by antimicrobial treatment. Changes in glycolipid ratios during treatment with both peptides suggests a modulation in the negative curvature stress of the membrane. Changes in surface curvature stress have been associated to increased resistance to antimicrobial peptide activity. Overall, lipidomic studies of pathogen response to the presence of antimicrobial agents can open avenues for research in investigating adaptability of microorganisms to these stress factors through changes in membrane composition.

## 6. Conflict of Interest

The authors declare that they have no known competing financial interests or personal relationships that could have appeared to influence the work reported in this paper.

## Acknowledgements

The research was funded by MinCiencias through the Grant Program (Cod. 1204-937-101846, CR 19576-2024). Additional support came from an internal grant from the Faculty of Sciences at Universidad de los Andes (INV-2025-213-3435). We would like to thank the Faculty of Sciences for granting the Academic Semester for Research (STAI semester 2026-1), which provided valuable time for completing this manuscript.

## Notes

### Competing Interest Statement

The authors have declared no competing interest.

